# Activation of α7 Nicotinic Acetylcholine Receptor Inhibits Neutrophil *Ex Vivo* Swarming and Pulmonary Recruitment in Inflamed Mice through an Integrin αMβ2-Dependent Mechanism

**DOI:** 10.64898/2026.10.06.752388

**Authors:** Jared L. Casteel, Tasha Phillips, Sanjay Singh, Tea Tsaava, Aisling Tynan, Valentin A. Pavlov, Alex Hopke, Valentin P. Yakubenko

## Abstract

**Background:** The cholinergic anti-inflammatory pathway regulates inflammation through vagus signaling that activates the α7 nicotinic acetylcholine receptor (α7nAChR) on leukocytes. Suppression of macrophage inflammatory cytokine production through α7nAChR-mediated inhibition of NF-κB activation is considered a central mechanism of this pathway. However, the effects of cholinergic activation of α7nAChR on neutrophils remain poorly understood. Here, we investigated whether and how activation of α7nAChR in neutrophils regulates neutrophil migration, defined the underlying molecular mechanism, and evaluated its contribution to anti-inflammatory protection during endotoxemia.

**Results:** The ex vivo swarming assays revealed that activation of α7nAChR with the selective agonists PNU-282987 and AR-R17779 significantly inhibited human and mouse neutrophil migration. The inhibitory effect of PNU-282987 was abolished in α7nAChR-deficient neutrophils, confirming receptor specificity. Mechanistically, α7nAChR activation reduced integrin α_M_β_2_ activation, decreased F-actin polymerization, and impaired neutrophil polarization. These effects required α_M_β_2_, as PNU-282987 failed to inhibit actin remodeling in α_M_-deficient neutrophils. Modulation of α_M_β_2_ activation with an activating antibody (MEM48) demonstrated that α7nAChR regulates neutrophil migration by limiting excessive α_M_β_2_ activation and maintaining an optimal adhesive state. *In vivo*, adoptively transferred fluorescently labeled PNU-282987-treated neutrophils exhibited reduced recruitment to the lungs compared with vehicle-treated neutrophils during LPS-induced endotoxemia. Furthermore, neutrophil depletion markedly diminished the protective effect of PNU-282987 on survival, demonstrating that neutrophils contribute to α7nAChR-mediated protection during systemic inflammation.

**Conclusions:** Our findings identify neutrophils as an important cellular target of the cholinergic anti-inflammatory pathway and demonstrate that α7nAChR activation limits neutrophil migration by suppressing integrin α_M_β_2_ activation and actin cytoskeletal remodeling. Together, these findings establish α_M_β_2_ as a critical downstream effector of α7nAChR signaling in neutrophils and identify regulation of integrin activation as a previously unrecognized mechanism of the cholinergic anti-inflammatory pathway.

## BACKGROUND

Inflammation is an essential host defense mechanism that protects against infection and tissue injury. Under physiological conditions, the inflammatory response is tightly regulated, localized, and resolved in a timely manner. Failure to restore immune homeostasis results in excessive or chronic inflammation that contributes to the pathogenesis of numerous diseases, including sepsis, rheumatoid arthritis, and cardiovascular disorders ^1–3^.

Over the past two decades, discoveries in neuroimmunology have established the vagus nerve as a critical regulator of inflammation and have provided a rationale for the development of bioelectronic and pharmacological anti-inflammatory therapies. In the inflammatory reflex, afferent vagal fibers sense inflammatory mediators generated during infection or tissue injury and relay this information to the brainstem, which coordinates efferent cholinergic signaling to suppress excessive immune activation. ^4, 5^ The efferent arm of this reflex, termed the cholinergic anti-inflammatory pathway, represents a major physiological mechanism that regulates inflammation in multiple organs, including the lungs, spleen, liver, and gastrointestinal tract ^6, 7^.

The principal receptor mediating cholinergic anti-inflammatory signaling is the α7 nicotinic acetylcholine receptor (α7nAChR), which is activated by endogenous acetylcholine and selective pharmacological agonists. Although originally characterized in neurons, α7nAChR is also expressed by numerous immune cells, including macrophages^8^, neutrophils ^9^, dendritic cells^10^, and lymphocytes^11^. Pioneering studies by Kevin Tracey and colleagues established macrophages as the primary cellular target of the cholinergic anti-inflammatory pathway by demonstrating that α7nAChR activation suppresses NF-κB signaling and inhibits the production of pro-inflammatory cytokines, including TNFα, IL-1β, and IL-6. ^8 12, 13^ This mechanism is recognized as a key outcome of cholinergic anti-inflammatory activation.

In contrast, the role of neutrophils as a target of the cholinergic anti-inflammatory pathway remains poorly understood. Studies have reported reduced neutrophil accumulation in inflamed tissues following α7nAChR activation during acute inflammation, yet the underlying mechanisms remain unclear.^9, 14, 15^ These effects have primarily been attributed to indirect regulation through endothelial α7nAChR signaling, altered vascular permeability, or changes in chemokine production. Whether α7nAChR directly regulates neutrophil migration and whether neutrophil modulation contributes to the protective effects of cholinergic stimulation during systemic inflammation remain unresolved.

In our recent publication, we revealed the contribution of α7nAChR to macrophage migration ^16, 17^. We demonstrated that the reduced survival of α7nAChR-deficient mice correlated with a decreased number of macrophages in the lungs during LPS-induced endotoxemia. Previous studies, including our own, have shown that recruitment of monocyte-derived macrophages to the lungs during experimental sepsis exerts a protective effect by accelerating neutrophil efferocytosis, thereby reducing pro-inflammatory stimulation^18, 19^. Our continued research demonstrated that α7nAChR deficiency impaired macrophage recruitment *in vivo* and *in vitro*^20^. Notably, analysis of macrophage adhesive receptors indicated that α7nAChR deficiency significantly reduced the expression of integrin α_M_β_2_ (Mac-1, CD11b/CD18), a β_2_ integrin expressed predominantly on myeloid leukocytes that regulates leukocyte adhesion, migration, phagocytosis, and inflammatory signaling. ^21^ Our results highlighted a potential role of α_M_β_2_ in the cholinergic response ^20^.

Guided by these findings, we hypothesized that the effects of α7nAChR activation on neutrophils are mediated through integrin α_M_β_2_. The role of α_M_β_2_ in inflammation is complex and context dependent, with studies demonstrating both detrimental ^22,21,23, 24^ and protective ^25,22^ functions depending on the disease model. Importantly, neutrophil activation by inflammatory mediators such as fMLP, LTB₄, LPS, and PMA rapidly mobilizes intracellular α_M_β_2_-containing granules, increasing surface expression of the receptor by approximately three- to fivefold within minutes^26^ ^23, 24^. This dynamic regulation places α_M_β_2_ at the center of neutrophil responses during acute inflammation, particularly in endotoxemia and sepsis.

In the present study, we investigated whether α7nAChR directly regulates neutrophil migration *ex vivo* and recruitment during endotoxemia *in vivo* and explored the molecular mechanism underlying this response. To model neutrophil migration *ex vivo*, we employed a neutrophil swarming assay using heat-killed Candida albicans. Neutrophil swarming is a coordinated migratory response that enhances antimicrobial defense but can also exacerbate tissue injury during uncontrolled inflammation^27, 28^. This process recapitulates key features of inflammatory neutrophil recruitment, including reactive oxygen species production, chemokine release, and coordinated cell migration, making it a physiologically relevant model for studying neutrophil inflammatory responses.

We demonstrate that cholinergic stimulation directly suppresses neutrophil swarming by limiting α_M_β_2_ activation, thereby attenuating actin cytoskeletal remodeling and reducing neutrophil recruitment to the lungs during LPS-induced systemic inflammation. These findings identify neutrophils as direct cellular targets of the cholinergic anti-inflammatory pathway and establish α_M_β_2_ as a previously unrecognized molecular effector linking cholinergic signaling to neutrophil migration.

## MATERIAL AND METHODS

### Reagents and Antibodies

The selective α7 nicotinic acetylcholine receptor (α7nAChR) agonist PNU-282987, lipopolysaccharide (LPS) from *Escherichia coli* O111:B4, poly-L-lysine solution (0.1%), polyvinyl alcohol (PVA), AR-R17779, PKH26, and fMLP were purchased from Sigma-Aldrich (St. Louis, MO, USA). Human peripheral blood fibrinogen was purchased from Enzyme Research Laboratories (South Bend, IN, USA). Recombinant laminin was obtained from R&D Systems (Minneapolis, MN, USA). Iscove’s Modified Dulbecco’s Medium (IMDM) was purchased from Corning (Corning, NY, USA). The cell-permeant fluorescent dye Calcein AM and mAb24 antibody was purchased from Invitrogen (Carlsbad, CA, USA). For immunofluorescence assays, Phalloidin 555 was obtained from Abcam (Cambridge, UK) and DAPI-enriched fluoroshield mounting medium was purchased from OriGene (Rockville, MD, USA). MEM48 antibody purchased from Satna Cruz Biotechnology (Dallas, TX, USA). EasySep Direct Human Neutrophil Isolation Kit was purchased from STEMCELL Technologies (Vancouver, BC, Canada), and the Neutrophil Isolation Kit (mouse) in conjunction with MACS LS columns purchased from Miltenyi Biotec (San Diego, CA, USA). anti-mouse Ly6G depletion antibody (clone 1A8) was purchased from BioLegend (San Diego, CA, USA), and the Alexa Fluor 647-conjugated anti-mouse/human integrin α_M_ antibody (clone M1/70) was purchased from BD Biosciences (San Jose, CA, USA;). Heat-killed *Candida albicans* targets were prepared in-house according to standard laboratory protocols.

### Animals

Wild-type C57BL/6J (stock #000664), integrin α_M_-deficient mice (B6.129S4-*Itgam^tm1Myd^*/J) and a7nAChR-deficient (a7nAChR^-/-^; B6.129S7-Chrna7tm1Bay/J, stock #003232) mice were purchased from the Jackson Laboratory (Bar Harbor, Maine). The integrin α_M_-deficient and α7nAChR-deficient lines were subsequently maintained as an in-house colony; homozygous knockout mice were generated by intercrossing heterozygous breeders and confirmed via tail-snip genotyping prior to experimental use. Both male and female mice aged 8–16 weeks were used across all experiments, as no sex-dependent differences in response to α7nAChR modulation were observed. Body weight was recorded for each animal and used to calculate weight-normalized injection volumes throughout. All animal procedures were approved by the East Tennessee State University Institutional Animal Care and Use Committee (IACUC) under protocol P240801-VA119 and conducted in accordance with its approved guidelines.

### LPS-Induced Endotoxemia Model

To evaluate the effect of α7nAChR activation on systemic inflammation and survival, an acute LPS-induced endotoxemia model was utilized as previously described.^29^ Wild-type C57BL/6J mice (n = 5 per group) were randomly assigned to treatment cohorts. Mice received a single intraperitoneal (IP) injection of either the selective α7nAChR agonist PNU-282987 (3 mg/kg) dissolved in DMSO (0.0625% v/v) or an equivalent volume of DMSO vehicle control. The concentration of PNU-282987 was selected based on our previous study ^29^, which demonstrated a protective effect at this concentration, and to maintain the final DMSO concentration in the injection solution below 0.1% to minimize the anti-inflammatory effect of DMSO. Following a 30-minute pretreatment period, endotoxemia was induced via IP injection of LPS (*Escherichia coli* O111:B4; Sigma-Aldrich) at a dose of 8 mg/kg body weight. Animals were monitored for clinical signs of endotoxemia and survival for up to 120 hours post-challenge.

### In Vivo Neutrophil Depletion Assay

To establish the specific contribution of neutrophils to α7nAChR-mediated protection during endotoxemia, in vivo neutrophil depletion was performed. Mice (n = 6 per group) were administered an IP injection of 100 µg of anti-mouse Ly6G monoclonal antibody (clone 1A8; BioLegend, San Diego, CA, USA) 24 hours prior to the induction of endotoxemia, followed by a second injection 24 hours post-LPS, a protocol previously validated to achieve robust peripheral neutrophil depletion at this dose^30^. Following a 30-minute pretreatment period, endotoxemia was induced via IP injection of LPS. In these experiments, the LPS dose was reduced by 25%, as the standard dose caused rapid and severe mortality in neutrophil-depleted mice. Animals were monitored for clinical signs of endotoxemia and survival for up to 96 hours post-challenge.

### Human and Mouse Neutrophil Isolation

The collection and isolation of human neutrophils were conducted in strict accordance with federal regulations and institutional guidelines, as previously described. All experimental protocols were reviewed and approved by the Institutional Biosafety and Chemical Safety Committee at East Tennessee State University. Fresh peripheral blood samples from healthy donors were purchased from a local BioIVT collection center. Blood was collected into EDTA tubes under protocols approved by the BioIVT Institutional Review Board, and all samples were de-identified before being provided for research use. To ensure maximum cell viability, human neutrophils were purified within 6 hours of collection using the negative-selection EasySep Direct Human Neutrophil Isolation Kit (STEMCELL Technologies) according to the manufacturer’s specified protocol.

For the isolation of non-activated mouse neutrophils, bone marrow was utilized. Briefly, both femurs and tibias were harvested from a single mouse per preparation and flushed with sterile PBS. Single-cell suspensions were prepared by passing the marrow through a 70 µm nylon mesh cell strainer. Mouse bone marrow neutrophils were subsequently purified via negative selection using the Neutrophil Isolation Kit (mouse) and MACS technology with LS columns according to the manufacturer’s instructions (Miltenyi Biotec, San Diego, CA, USA).

### Ex Vivo Neutrophil Swarming Assay

Neutrophil swarming was investigated using a modified micro-patterning approach ^31, 32^. Briefly, a microarray printing platform (Picospotter; PolyPico) was utilized to microprint 0.1% poly-L-lysine (Sigma-Aldrich) onto glass slides (Fisher Scientific) to form an 8×8 grid, with individual spots approximately 100 µm in diameter (Figure 2A). Slides were dried at room temperature for approximately 3 hours. Heat-killed *Candida albicans* was prepared by incubation at 70°C for 20 minutes and used as the swarming target. The dead microorganisms initiate <u>neutrophil swarming</u> but exclude signaling, which may influence neutrophil migration. A suspension of heat-killed *C. albicans* was added to each well of a 16-well ProPlate slide chamber system (Grace Bio-Labs) pre-attached to the patterned glass slides and incubated with rocking for 5–10 minutes. Unbound *C. albicans* targets were removed by thorough washing with PBS.

Following target immobilization, slides were coated with 10 µg/mL of fibrinogen for 3 hours at 37°C and subsequently washed three times with PBS. Human peripheral blood neutrophils or mouse bone marrow neutrophils (isolated via negative selection as described above) were fluorescently labeled with 10 µM Calcein AM (Invitrogen) for 30 minutes at 37°C per manufacturer’s protocol or using PKH26 according to manufacturer’s protocol. Following labeling, cells were pelleted by centrifugation and resuspended in Iscove’s Modified Dulbecco’s Medium (IMDM). Prior to plating, cells were pretreated with either varying concentrations of fMLP (0, 1, 10, or 100 nM) for 15 minutes, or with 30 µM PNU-282987, 10 µM AR-R17779, at 37°C, or an equivalent concentration of DMSO vehicle control for 30 minutes at 37°C. The concentration of PNU-282987 was selected based on the range reported in published studies (10–150 µM) ^33, 34^ and our preliminary experiments evaluating different concentrations of PNU-282987 and its vehicle, DMSO, in neutrophil swarming assays. DMSO affected neutrophil swarming at concentrations above 0.125%, which corresponds to the DMSO concentration required to prepare 60 µM PNU-282987. Therefore, we selected 30 µM PNU-282987 for our experiments to minimize potential vehicle-related effects. The biological activity of PNU-282987 at this concentration was also demonstrated in our previous publication ^29^. The concentration of the water-soluble α7nAChR agonist AR-R17779 (10 µM) was selected from the range (10-100 µM) based on previously published studies ^35 36^.

For the alteration of integrin α_M_β_2_ activation, cells were treated with 10 μg/mL of the integrin β_2_-activating monoclonal antibody MEM-48 (Santa Cruz Biotechnology) for 15 minutes at 37°C after incubation with PNU-282987 as described above.

Treated cells were seeded onto prepared slides at a final concentration of 1 x 10⁶ cells per mL. Each well was imaged using an 4x objective on an EVOS FL Auto Microscope (Invitrogen) equipped with an environmental humidity chamber at 37°C, with time-lapse images captured every 2 minutes for a total of 2 hours. Swarming behavior was quantified by measuring the mean fluorescent intensity around each target spot over time using ImageJ software.

### Phalloidin Immunofluorescence Staining and Quantification of Neutrophil Swarming

To evaluate F-actin distribution and neutrophil morphology, immunofluorescence staining was performed on static adherent neutrophils and swarming neutrophil clusters as previously described with modifications^37^. Assays were executed on either blank glass slides (static adhesion) or micro-patterned slides featuring *Candida albicans* targets (swarming). Calcein AM-labeled human or mouse neutrophils were pretreated with 30 µM PNU-282987 or DMSO vehicle control for 30 minutes at 37°C before addition to the slides. For static adhesion assays on blank slides, cells were allowed to adhere for 30 minutes at 37°C in a humidified 5% CO₂ incubator, after which non-adherent cells were gently aspirated and remaining adherent cells were fixed with 2% PFA. Swarming assays were terminated 45 minutes post-initiation by adding 16% paraformaldehyde directly into the wells to achieve a final concentration of 2% PFA. All preparations were fixed for 30 minutes in the dark at room temperature, washed three times with PBS, and permeabilized with 0.1% Triton X-100 in PBS for 5 minutes at room temperature. Following three additional PBS washes, cells were incubated overnight at 4°C in a humidified chamber with a staining cocktail containing Phalloidin 555 (1:1,000; Abcam) and Alexa Fluor 647-conjugated anti-CD11b (clone M1/70; 1:200 in 1% BSA/PBS; BD Biosciences). Following three gentle PBS rinses, swarming slides were mounted with DAPI-enriched mounting media (OriGene) to label cell nuclei and sealed.

Fluorescent images were acquired on a Leica TCS SP8 confocal microscope. Static adherent cells were captured using a 20× objective to evaluate cell density and integrated intensity across broader fields of view. Swarming clusters were imaged using a 63× oil immersion objective for detailed morphological resolution, and raw files were exported using Leica LAS X software. Quantitative image analysis was conducted using a custom CellProfiler pipeline^38^. For swarming assays, individual cell nuclei were identified from the DAPI channel using adaptive Otsu thresholding with intensity-based declumping, and cell boundaries were defined from the phalloidin channel using a propagation algorithm (regularization factor 0.05) seeded from nuclear masks; objects touching image borders or outside a defined size range were excluded. F-actin polymerization across both conditions was determined by measuring the integrated fluorescent intensity of the Phalloidin 555 channel within cellular boundaries. Data were normalized to the median of each of three independent experiments, and statistical outliers were identified and removed using the ROUT method.

### Static Cell Adhesion Assay

Cell adhesion was evaluated using a modified protocol based on established fluorescent labeling methods ^37, 39, 40^. Immulon 4HBX 96-well plates (Thermo Fisher Scientific) were coated with 10 μg/mL of human fibrinogen (Enzyme Research Laboratories) for 3 hours at 37°C. Wells were subsequently blocked with 0.5% polyvinyl alcohol (PVA) for 1 hour at room temperature. Primary human or mouse bone marrow-derived neutrophils were fluorescently labeled with 10 μM Calcein AM (Invitrogen) for 30 minutes at 37°C, washed with PBS, and resuspended in HBSS to a final concentration of 1 x 10^6^. Prior to plating, labeled cells were pretreated with either 30 μM PNU-282987 or an equivalent volume of DMSO vehicle control for 30 minutes at 37°C. Aliquots of the treated cell suspensions (5 x 10^4^ cells/well in 50μL) were added to the prepared plates and incubated for 30 minutes at 37°C in a humidified 5% CO_2_ incubator. Total initial fluorescence was measured using an Agilent BioTek Synergy H1 fluorescence plate reader (Agilent BioTek, Santa Clara, CA, USA) to establish baseline cell input. Non-adherent cells were removed by washing with sterile PBS, and the remaining adherent cell fluorescence was quantified. The percentage of adherent cells was calculated by comparing the post-wash fluorescent signal against the corresponding pre-wash baseline measurement.

### Phalloidin Immunofluorescence Staining of Adherent Neutrophils

To evaluate F-actin distribution in static adherent cells, neutrophils were prepared on unmodified glass slides coated directly with fibrinogen, matching the surface chemistry of the swarming assays but lacking both poly-L-lysine micro-patterning and Candida albicans targets. Slides were coated with 10 µg/mL of fibrinogen for 3 hours at 37°C. Calcein AM-labeled human or mouse neutrophils were pretreated with 30 µM PNU-282987 or DMSO vehicle control for 30 minutes at 37°C before being seeded onto the coated slides. Cells were allowed to adhere for 30 minutes at 37°C in a humidified 5% CO_2_ incubator. Following incubation, media containing non-adherent cells was gently aspirated and the remaining adherent cells were immediately fixed by addition of 4% PFA for 30 minutes in the dark at room temperature. Cells were washed three times with PBS and permeabilized using 0.1% Triton X-100 in PBS for 5 minutes at room temperature. Following three additional PBS washes, cells were incubated overnight at 4°C in a humidified chamber with the staining cocktail containing Phalloidin 555 (1:1,000; Abcam) and Alexa Fluor 647-conjugated anti-α_M_ (clone M1/70; 1:200; BD Biosciences). Fluorescent images were captured using a Leica TCS SP8 confocal microscope equipped with a 20x objective to allow quantification of adherent cell numbers across broader fields of view, and downstream integrated fluorescent intensity quantification of the actin channel was performed via CellProfiler as described above.

### Flow Cytometric Analysis of integrin α_M_ Expression and Activation

To evaluate total and active integrin α_M_ surface expression, primary human neutrophils (1 x 10^6^ cells/mL) suspended in HBSS were treated with varying concentrations of fMLP (0, 1, 10 or 100 nM) for 15 minutes at 37°C. Following fMLP treatment, cells were simultaneously incubated with either an Alexa Fluor 647-conjugated anti-human integrin α_M_ antibody (clone M1/70; BD Biosciences) and a PE-conjugated conformation-specific anti-active integrin β_2_ antibody (clone mAb24; BD Biosciences) for 30 minutes at 37°C to monitor receptor translocation and conformational affinity shifts during stimulation. Following incubation, cell suspensions were immediately fixed by the addition of paraformaldehyde to a final concentration of 2% PFA for 15 minutes to stabilize surface antibody complexes. Fixed cells were washed twice with cold stain buffer (PBS containing 1% BSA), pelleted at 300 rcf, and resuspended in PBS. Fluorescence intensity was acquired on a BD Fortessa X-20 flow cytometer. Data analysis was conducted using FACSDiva software, and results are expressed as mean fluorescence intensity (MFI) relative to the untreated control group from each experimental day.

### In-Vivo Adoptive Transfer Assay

Primary mouse neutrophils were isolated from the bone marrow of two wild-type C57BL/6J donor mice per experiment as described above. Purified donor neutrophils were fluorescently labeled with either PKH67 (green) or PKH26 (red) cell linker kits (Sigma-Aldrich) according to the manufacturer’s instructions. Based on our previous studies^29, 41, 42^, 2 μL of PKH67 or 1 μL of PKH26 per 1 × 10⁶ cells was used to achieve comparable fluorescence intensities for in vivo cell tracking. We previously demonstrated that reciprocal labeling with PKH67 and PKH26 yields equivalent migration results, indicating that the observed cell migration is independent of the fluorescent dye used for labeling^29^. Labeled cells (1 × 10⁶ cells/mL) were treated ex vivo with either 30 μM PNU-282987 or an equivalent volume of DMSO vehicle control for 30 minutes at 37°C. Following treatment, cells were washed and counted. Equal numbers of PNU-treated and vehicle-treated neutrophils (0.75-1.5 × 10⁶ per group) were mixed together in sterile PBS and administered via tail vein injection (100µL) into recipient C57BL/6J mice. Recipient mice were immediately challenged via intraperitoneal injection with a sublethal dose of LPS (8 mg/kg; Escherichia coli O111:B4; Sigma-Aldrich). At 24 hours post-injection, mice were humanely euthanized and systemic circulation was cleared by transcardial perfusion with cold PBS. Lungs were harvested and enzymatically digested as previously described^29^. Single-cell suspensions were recovered and the relative recruitment of PKH67- and PKH26-labeled neutrophils in digested lung tissue was quantified by flow cytometry on a BD Fortessa X-20 flow cytometer and analyzed by FACSDiva software.

### Statistical Analysis

Statistical analyses were performed using GraphPad Prism 11 (GraphPad Software, Boston, MA, USA). Continuous data were first evaluated for normality; the Shapiro-Wilk test was applied for smaller sample sizes and the D’Agostino-Pearson test was used for larger cohorts, with *α* = 0.05 as the threshold for normality. For comparisons between two independent groups, a two-tailed Student’s t-test was utilized for normally distributed data, and a Mann-Whitney U test was applied as the non-parametric alternative for datasets that consistently failed normality criteria following logarithmic or alternative transformations. For comparisons among three or more independent groups, data were analyzed using a one-way non-parametric ANOVA (Kruskal-Wallis test) followed by Dunn’s multiple comparisons post-hoc test. However, to prevent artificial donor clustering in assays evaluating heterogeneous single-cell or spot-level behaviors (such as micropatterned swarming and immunofluorescence imaging), pooled data from these experiments were strictly analyzed using unpaired tests (e.g., Mann-Whitney U test) to treat individual spots and cells as independent micro-environments. Longitudinal survival data over the 120-hour monitoring window were analyzed using the Log-rank (Mantel-Cox) test. Quantitative data are presented according to experimental design: wide-field and confocal single-cell imaging datasets are presented as individual cellular data points displaying the median bar, whereas other endpoint assays are expressed as the mean ± standard error of the mean (SEM). For all analyses, p<0.05 was considered statistically significant.

## RESULTS

### Activation of α7nAChR inhibits ex vivo neutrophil swarming to *C. albicans*

To determine whether activation of α7nAChR directly regulates neutrophil migration, we performed an ex vivo neutrophil swarming assay. Heat-killed *C. albicans* was immobilized on printed spots of ProPlate wells (Grace Bio-Labs) to serve as swarming targets (Fig. 1A). Wells were coated with fibrinogen (10 µg/mL), a physiological ligand for integrin α_M_β_2_ that supports neutrophil migration. First, we tested human neutrophils, isolated from peripheral blood. Purified neutrophils were labeled with the fluorescent dye PKH26, and pretreated with the selective α7nAChR agonist PNU-282987 (30 μM) or vehicle for 30 min before the assay. A total of 2 × 10⁵ neutrophils were added to each well, and swarm formation was monitored for 2 h using a fully automated EVOS imaging system equipped with temperature and CO₂ control for live-cell time-lapse microscopy.

**Figure 1.**
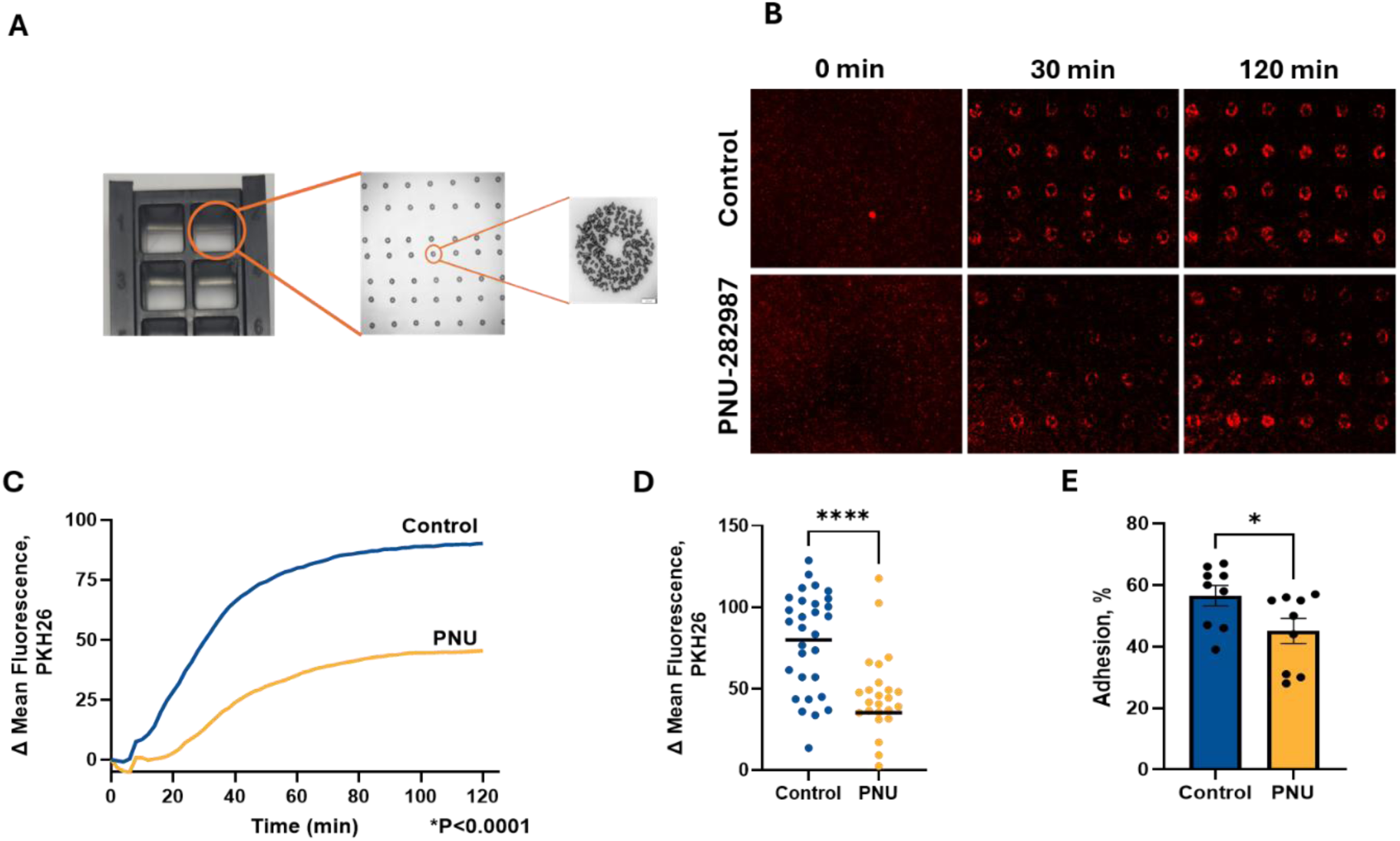
α7nAChR activation reduces human neutrophil swarming and static adhesion. **(A)** Schematic of the micro-patterned swarming assay: glass slides were microprinted with an 8×8 grid of poly-L-lysine spots (∼100 µm diameter) using a Picospotter platform, coated with heat-killed *Candida albicans*, and assembled into a 16-well ProPlate chamber system. **(B)** Freshly isolated human neutrophils were stained with PKH26 and added to wells pre-coated with 10 µg/mL fibrinogen and treated with 30 µM PNU-282987 or vehicle and swarming toward *C. albicans* targets was imaged at 4x magnification by time-lapse microscopy every 2 min for 2 h; representative images are shown. **(C)** Change in mean fluorescence intensity (ΔMFI, PKH26) at swarm spots, calculated relative to baseline (*t* = 0) intensity for each spot, was quantified over the 2 h imaging period; data represent the mean of all spots pooled from 3 independent experiments, each performed on an individual donor (8–10 spots/experiment/condition). **(D)** ΔMFI at the 60-min time point is shown for all individual spots pooled from the 3 independent experiments in (C), with the line representing the mean; data were log-transformed to meet normality assumptions and analyzed by Welch’s t-test. ****p<0.0001. **(E)** Fibrinogen (10 µg/mL) was immobilized on 96-well plates for 3 h at 37°C. Calcein-AM-labeled human neutrophils were pretreated for 30 min with vehicle or PNU-282987 (30 µM), added to fibrinogen-coated wells, and allowed to adhere for 30 min; adhesion was quantified by fluorescence plate reader. All 9 data points from 3 independent experiments (one donor per experiment) are shown as mean ± SEM; data were analyzed by Mann-Whitney U test. **p<0.01.

Control neutrophils (vehicle-treated) rapidly migrated toward *C. albicans*, forming dense swarms within 30–40 min. In contrast, pretreatment with PNU-282987 significantly reduced neutrophil swarming throughout the assay (Fig. 1B–D, Supplemental Video 1), indicating that activation of α7nAChR suppresses neutrophil migratory responses. Since adhesion is an essential step of cell migration, we tested the effect of α7nAChR activation on neutrophil adhesion to fibrinogen. We detected a statistically significant decrease in adhesion of PNU-282987 – treated neutrophils (Fig. 1E).

To confirm that this effect was mediated by α7nAChR rather than by an off-target action of PNU-282987, neutrophils were treated with another selective α7nAChR agonist, AR-R17779 (10 μM), for 30 min before the assay. Similar to PNU-282987, AR-R17779 significantly inhibited neutrophil swarming (Fig. 2A, B, Supplemental Video 2), further supporting a role for α7nAChR signaling in regulating neutrophil migration.

**Figure 2.**
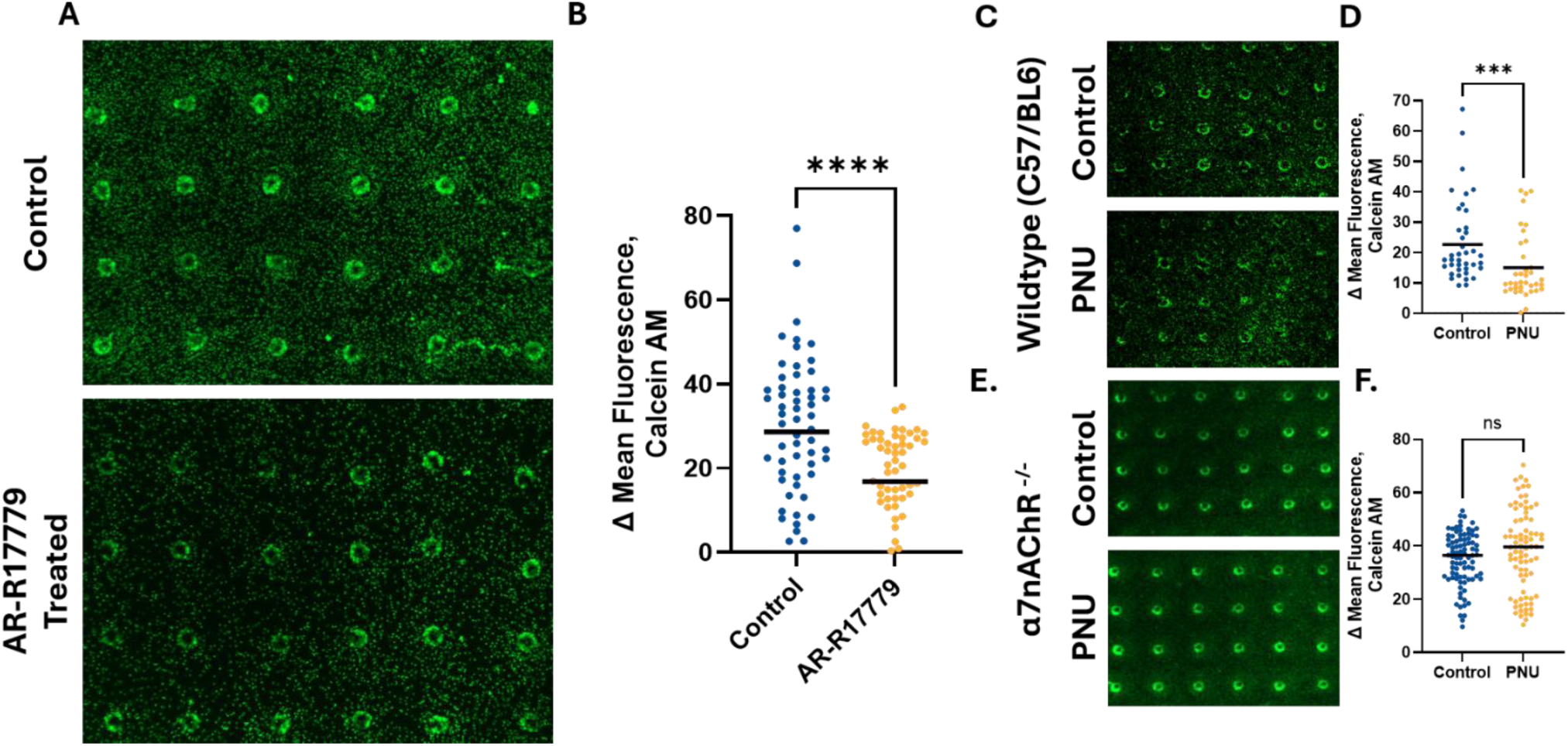
α7nAChR activation decreases human neutrophil swarming with an alternative agonist (AR-R17779) and reduces swarming in WT, but not α7nAChR^-/-^, mouse neutrophils. **(A)** Representative microscopic image (4x magnification) at the 60-min timepoint of human neutrophil swarming performed as in Figure 1, using AR-R17779 (10 µM) in place of PNU-282987. **(B)** Change in mean fluorescence intensity (ΔMFI, calcein AM) at the 60-min timepoint is shown for all individual spots (10–20 spots/experiment) pooled from 3 independent experiments (one donor per experiment), with the line representing the mean; data failed normality testing and were analyzed by Mann-Whitney U test. ****p < 0.0001. **(C,E)** Bone marrow neutrophils were isolated from WT (C) or α7nAChR^-/-^ (E) mice using a Miltenyi Biotec MACS kit and assayed for swarming on fibrinogen (10 µg/mL) as in Figure 1, with cells treated with PNU-282987 (30 µM) or vehicle control. Representative microscopic image of WT neutrophil swarming at the 60-min timepoint is shown. **(D)** ΔMFI at the 60-min timepoint for WT neutrophils from (C) is shown for all individual spots pooled from 3 independent experiments (one mouse per experiment), with the line representing the mean; data failed normality testing and were analyzed by Mann-Whitney U test. ***p < 0.001. (F) ΔMFI at the 60-min timepoint for α7nAChR^-/-^ neutrophils **(E)** is shown for all individual spots pooled from 3 independent experiments (one mouse per experiment), with the line representing the mean; data were analyzed by Mann-Whitney U test. ns, p = 0.1152.

To further establish the requirement for α7nAChR, we examined swarming of neutrophils isolated from α7nAChR-deficient mice. Initially, we confirmed that neutrophils from wild-type mice exhibited an effect comparable to that of human neutrophils and that PNU-282987 markedly suppressed swarm formation (Fig. 2C, D, Supplemental Video 3). In contrast, although α7nAChR^-/-^ neutrophils retained robust swarming activity, treatment with PNU-282987 failed to inhibit their migration (Fig. 2E, F, Supplemental Video 4). These findings confirm that the inhibitory effect of PNU-282987 on neutrophil swarming is dependent on α7nAChR expression.

### Activation of α7nAChR suppresses actin polymerization during neutrophil migration

Because directed neutrophil migration requires coordinated actin remodeling, we next examined whether α7nAChR activation alters cytoskeletal organization during neutrophil swarming. Neutrophils were allowed to swarm toward immobilized *C. albicans* for 45 min, immediately fixed, and stained for F-actin, integrin α_M_β_2_, and nuclei. Integrin α_M_β_2_ was analyzed because it is strongly upregulated during inflammation and plays a critical role in neutrophil swarming.^37^

Confocal microscopy revealed that vehicle-treated neutrophils exhibited the characteristic polarized migratory phenotype, with prominent F-actin accumulation (green) at the leading edge and formation of a trailing uropod (Fig. 3A, B). In contrast, neutrophils treated with PNU-282987 displayed markedly reduced actin polymerization and impaired cell polarization, consistent with their diminished swarming activity. Notably, PNU-282987 failed to alter actin organization in α7nAChR-deficient neutrophils (Fig. 3C, D), demonstrating that inhibition of actin remodeling requires direct or indirect α7nAChR signaling.

**Figure 3.**
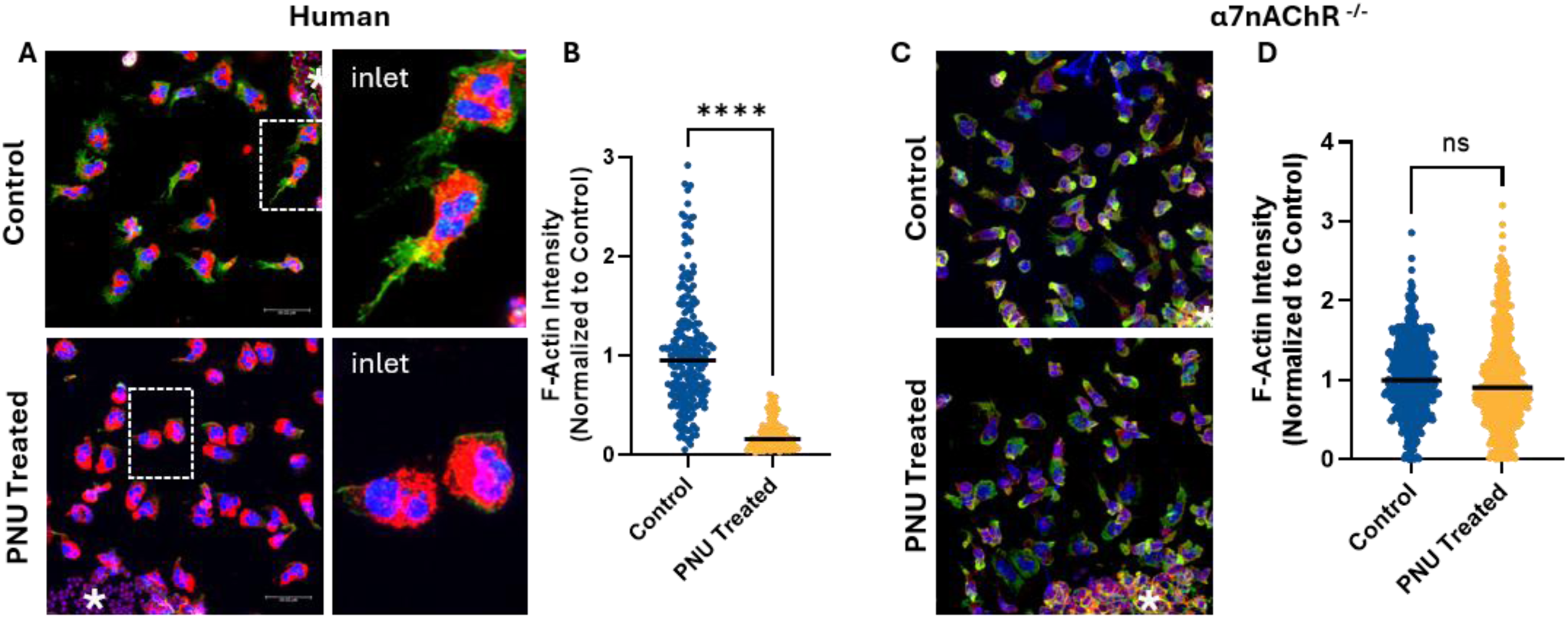
α7nAChR activation reduces F-actin polymerization during neutrophil swarming. **(A)** Human neutrophils were allowed to swarm on micropatterned slides coated with 10 µg/mL fibrinogen in the presence of PNU-282987 (30 µM) or vehicle control. At 45 min, cells were fixed with 4% PFA, permeabilized with 0.1% Triton X-100, and stained with DAPI, AF647-conjugated anti-αM, and Phalloidin-555, then imaged by confocal microscopy at 63x (oil immersion). Representative images are shown; asterisks in the corners indicate the position of the microprinted *C. albicans* spot. **(B)** F-actin intensity per cell from (A) was quantified using CellProfiler as described in materials and methods section. Mean F-actin intensity per cell (total phalloidin pixel intensity within the cell mask, divided by mask area) was calculated for each cell, normalized to the median of each of 3 independent experiments (one donor per experiment), and outliers were removed using the ROUT method. Data are shown for all individual cells, with the line representing the median; groups were compared by Mann-Whitney U test. ****p < 0.0001. **(C)** α7nAChR^-/-^ mouse bone marrow neutrophils were treated and imaged as in (A). Representative images are shown; asterisks in the corners indicate the position of the microprinted *C. albicans* spot. **(D)** F-actin intensity of α7nAChR^-/-^ mouse bone marrow neutrophils treated and imaged as in (B) was quantified and analyzed as in (C) using Mann-Whitney U test. ns, p = 0.4553.

### Integrin α_M_β_2_ is required for α7nAChR-mediated regulation of actin polymerization and neutrophil recruitment

Actin polymerization is initiated by integrin-mediated outside-in signaling following ligand engagement. Because integrin α_M_β_2_ is the principal neutrophil receptor regulating adhesion, migration, and outside-in signaling, we investigated whether α7nAChR-mediated changes in actin remodeling depend on α_M_β_2_ by comparing F-actin formation in WT and αM-deficient neutrophils following adhesion to fibrinogen.

Human and mouse neutrophils were pretreated with PNU-282987 or vehicle and allowed to adhere to fibrinogen for 30 min before fixation and staining. Consistent with the swarming assay, α7nAChR activation markedly reduced F-actin formation in adherent neutrophils from both species (Fig. 4 A-D), indicating that α7nAChR suppresses actin cytoskeletal remodeling independently of the swarming assay. In contrast, PNU-282987 failed to reduce actin polymerization in α_M_-deficient neutrophils, demonstrating that α_M_β_2_ is essential for α7nAChR-mediated regulation of actin remodeling (Fig. 4 E, F).

**Figure 4.**
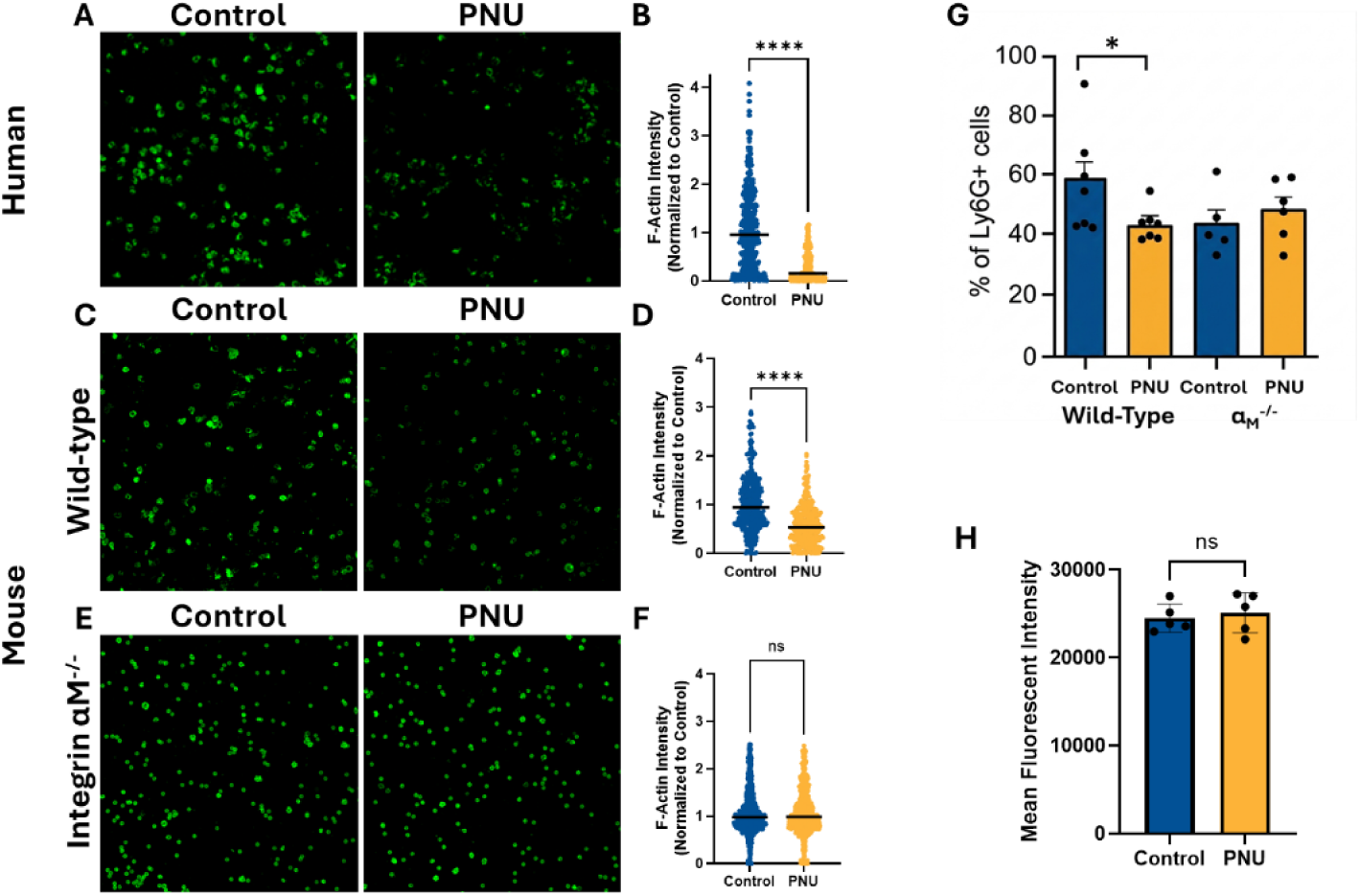
α7nAChR activation decreases actin polymerization during static adhesion in human and WT mouse neutrophils and requires integrin αMβ2. **(A)** Representative confocal images (20x) of phalloidin staining of human neutrophils following static adhesion to immobilized fibrinogen (10 µg/mL) for 30 min under Control (vehicle) or PNU-282987 (30 µM) treatment. **(B)** F-actin intensity per cell from (A) was quantified and normalized to Control; data were analyzed by Mann-Whitney U test. ****p < 0.0001. **(C)** Representative confocal images (20x) of phalloidin staining of wild-type (WT) mouse bone marrow-derived neutrophils following static adhesion to fibrinogen under Control or PNU-282987 treatment as in (A). **(D)** F-actin intensity per cell from (C) was quantified and normalized to Control; data were analyzed by Mann-Whitney U test. ****p < 0.0001. **(E)** Representative confocal images (20x) of phalloidin staining of integrin αM^-/-^ mouse bone marrow-derived neutrophils following static adhesion to fibrinogen under Control or PNU-282987 treatment as in (A). **(F)** F-actin intensity per cell from (E) was quantified and normalized to Control; data were analyzed by Mann-Witney test. n.s. p=0.871. **(G)** WT and αM^-/-^ mice were injected with 3mg/kg PNU-282987 or vehicle, followed by a sublethal dose of LPS 15 mins later. After 48h, lungs were removed, digested and analyzed using FACS. Neutrophils were identified as CD11b⁺/Ly6G⁺/F4/80⁻ cells or CD11c⁺/Ly6G⁺/F4/80⁻ for αM-deficient mice. Plot depicts the percentage of Ly6G^+^ cells. Data points represent individual mice with mean ± SEM (n=5-7 mice/group); analyzed via Welch’s t-test. *p<0.05. **(H)** The level of integrin αMβ2 expression (Mean Fluorescent Intensity) was detected on Ly6G^+^ cells collected from the digested lungs of WT mice at 48 hours after LPS administration. Data points represent individual mice with mean ± SEM (n=5 mice/per group). Data were analyzed by Welch’s t-test, n.s. p=0.652.

To determine whether α_M_β_2_ is also required for α7nAChR-mediated regulation of neutrophil recruitment *in vivo*, we used the mouse model of LPS-induced endotoxemia. In our previous study, we demonstrated that PNU-282987 treatment reduces neutrophil accumulation in the lungs of WT mice following LPS challenge ^29^. Here, we examined whether this response is altered in α_M_-deficient mice. WT and α_M_-deficient mice were pretreated with PNU-282987 or vehicle before administration of a sublethal dose of LPS. After 48 h, mice were perfused, lungs were collected and enzymatically digested, and neutrophils were quantified by flow cytometry as CD11b⁺/Ly6G⁺/F4/80⁻ cells. To account for the absence of the CD11b subunit in αM-deficient mice, lung neutrophils in this cohort were alternatively gated as CD11c⁺/Ly6G⁺/F4/80⁻ cells.

While PNU-282987 significantly reduced neutrophil recruitment to the lungs of WT mice, this effect was completely abolished in α_M_-deficient mice (Fig. 4G), demonstrating that α_M_β_2_ is required for α7nAChR-mediated regulation of neutrophil recruitment during endotoxemia.

To determine whether the reduced neutrophil recruitment resulted from altered α_M_β_2_ expression, we next measured surface α_M_β_2_ levels on lung neutrophils from WT mice treated with PNU-282987 or vehicle following LPS challenge. Surprisingly, PNU-282987 did not alter α_M_β_2_ surface expression (Fig. 4H). Because α_M_β_2_ was required for the inhibitory effect of α7nAChR despite unchanged receptor expression, we hypothesized that cholinergic stimulation regulates α_M_β_2_ activation rather than its surface expression.

### α7nAChR regulates neutrophil swarming through modulation of integrin α_M_β_2_-mediated adhesion

To investigate the potential contribution of α7nAChR to the regulation of α_M_β_2_ activation, neutrophils were pretreated with the α_M_β_2_-activating antibody MEM48 before the swarming assay. As expected, activation of α_M_β_2_ significantly inhibited neutrophil swarming, consistent with the theory of mesenchymal cell migration, which postulates that very strong adhesion inhibits cell motility (as described below in the Discussion section). Notably, pretreatment with PNU-282987 largely restored neutrophil migration despite the presence of the activating antibody, suggesting that α7nAChR signaling counteracts excessive α_M_β_2_ activation (Fig. 5A, B,C, Supplemental Video 5).

**Figure 5.**
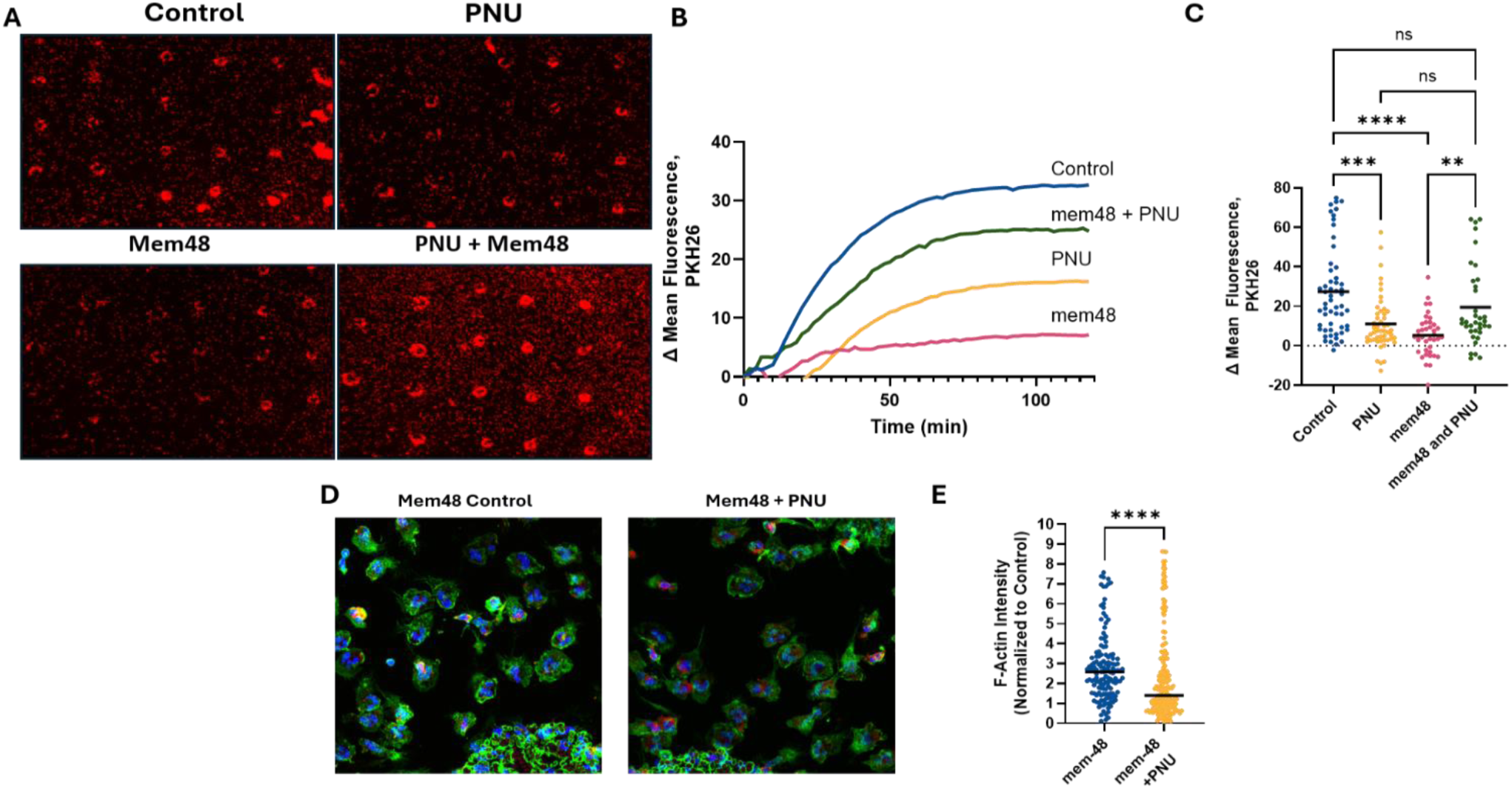
α7nAChR activation reduces αMβ2 integrin affinity and neutrophil migration during swarming. Human neutrophils were pretreated for 30 min with vehicle control, PNU-282987 (30 µM), the αMβ2-activating antibody Mem48 (10 µg/mL), or a combination of PNU-282987 and Mem48, prior to swarming against heat-killed *C. albicans* targets on slides coated with 10 µg/mL fibrinogen. **(A)** Representative swarming images for each treatment condition. **(B)** Kinetic quantification of neutrophil swarm size (ΔMFI, PKH26) over time; data represent the mean of 10–20 spots/experiment pooled from 3 independent experiments (one donor per experiment). **(C)** Swarm size (ΔMFI) at the 60-min timepoint for all treatment conditions; data were analyzed by one-way non-parametric ANOVA (Kruskal-Wallis) with Dunn’s multiple comparisons test. Control vs. PNU-282987, ***p < 0.001; Control vs. Mem48, ****p < 0.0001; Mem48 vs. Mem48+PNU-282987, **p < 0.01; PNU-282987 vs. Mem48+PNU-282987, ns; Control vs. Mem48+PNU-282987, ns. **(D)** Human neutrophils treated with Mem48 with or without PNU-282987 were fixed, permeabilized, and stained with DAPI, AF647-conjugated anti-integrin αM, and Phalloidin-555 as in Figure 3, and imaged by confocal microscopy at 63x (oil immersion). Representative images are shown. **(E)** F-actin intensity per cell was quantified as in Figure 3 and compared by Mann-Whitney U test. ****p < 0.0001.

Analysis of actin organization further supported this mechanism. Neutrophils treated with MEM48 exhibited intense cortical F-actin accumulation, forming a prominent actin ring characteristic of highly adherent cells with limited migratory capacity. In contrast, addition of PNU-282987 markedly reduced F-actin polymerization and improved front-rear polarity, consistent with the recovery of neutrophil swarming (Fig. 5D, E). These findings indicate that activation of α7nAChR reverses the cytoskeletal changes induced by excessive α_M_β_2_ activation.

### The degree of integrin α_M_β_2_ activation regulates neutrophil swarming

Because leukocyte migration is critically dependent on the degree of integrin activation, we next examined how different levels of α_M_β_2_ activation affect neutrophil swarming. Neutrophils were stimulated with increasing concentrations of fMLP (1–100 nM) for 15 min before the assay. Stimulation with 1 nM fMLP modestly enhanced neutrophil swarming, whereas higher fMLP concentrations progressively reduced swarming and completely inhibited neutrophil motility at 100 nM (Fig. 6A, B, C, Supplemental Video 6). Flow cytometric analysis confirmed that fMLP induced a concentration-dependent increase in both surface α_M_β_2_ expression and the high-affinity conformation of the receptor (Fig. 6D, E). Pretreatment with 1 nM fMLP increased α_M_β_2_ activation by approximately twofold and produced a modest enhancement of neutrophil swarming. In contrast, 10 nM fMLP increased total α_M_β_2_ expression by approximately twofold and the high-affinity conformation by nearly fivefold, resulting in a significant reduction in migration. Stimulation with 100 nM fMLP further enhanced α_M_β_2_ activation and almost completely suppressed neutrophil swarming. Together, these findings demonstrate that neutrophil migration is optimal at a limited level of α_M_β_2_ activation. Low activation and weak adhesion slow down migration, whereas excessive integrin activation promotes strong adhesion and restricts cell motility.

**Figure 6.**
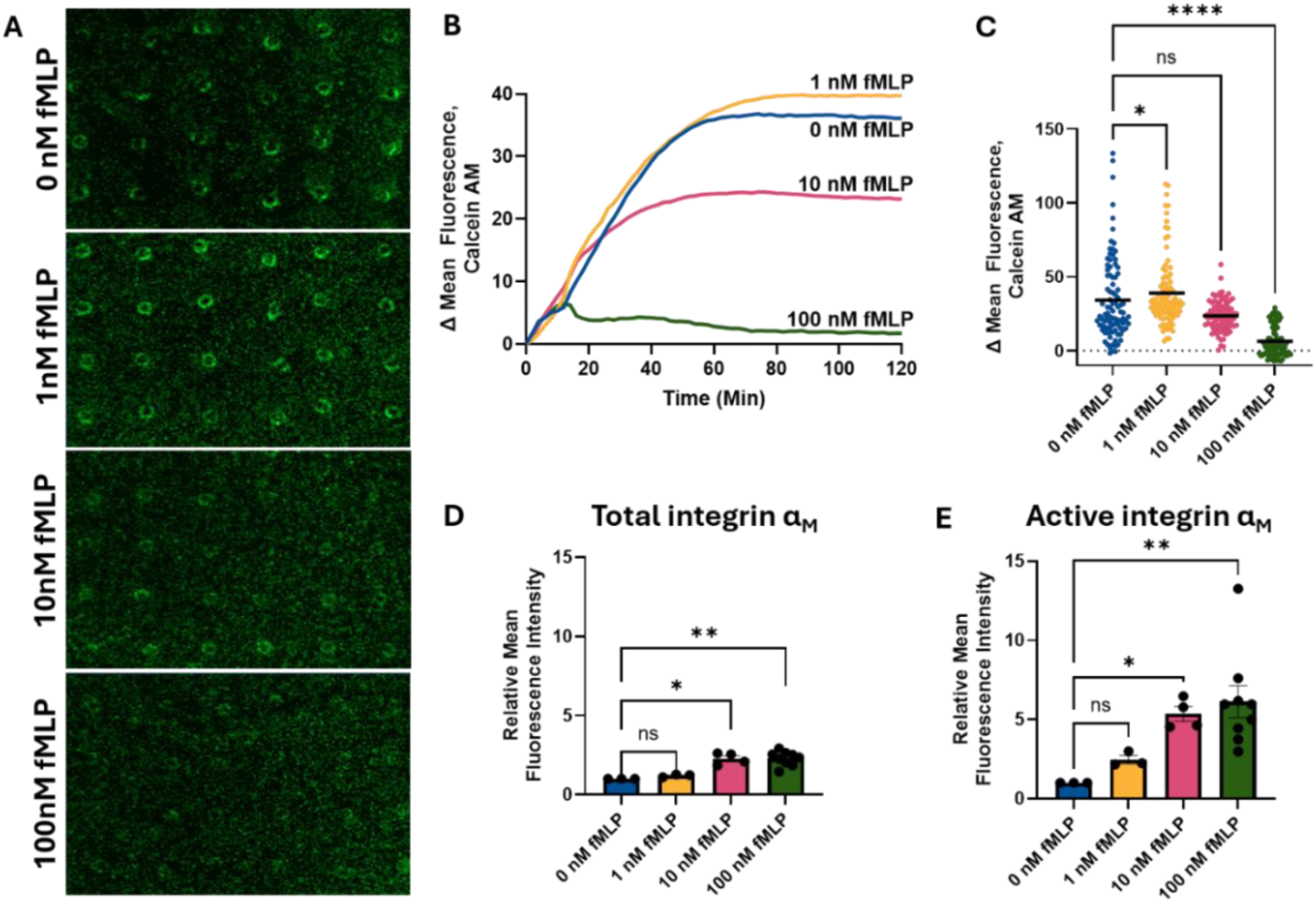
fMLP-treated neutrophils exhibit reduced swarming migration due to increased αM activation. **(A)** Human neutrophils were loaded onto swarming wells coated with 10 µg/mL fibrinogen following a 15-min pretreatment with 0, 1, 10, or 100 nM fMLP, and swarming against heat-killed *Candida albicans* targets was imaged by time-lapse microscopy at 4x magnification; representative 60-min timepoint images of 3 independent experiments are shown (one donor per experiment). **(B)** Kinetic quantification of swarm size (ΔMFI) over 120 min; data represent the mean of 10–20 spots/experiment pooled from 3 independent experiments. **(C)** Swarm size (ΔMFI, calcein AM) at the 60-min timepoint is shown for all individual spots, with the line representing the mean; data were analyzed by one-way non-parametric ANOVA (Kruskal-Wallis) with Dunn’s multiple comparisons test against the 0 nM fMLP control. 0 vs. 1 nM, *p < 0.05; 0 vs. 10 nM, ns; 0 vs. 100 nM, ****p < 0.0001. **(D)** Total αM surface expression was measured by flow cytometry following 15 minute treatment with 0, 1, 10, or 100 nM fMLP; data are shown as relative mean fluorescence intensity (MFI) normalized to the untreated control from each experimental day and were analyzed by one-way non-parametric ANOVA with Dunn’s multiple comparisons test against control. 0 vs. 1 nM, ns; 0 vs. 10 nM, *p < 0.05; 0 vs. 100 nM, **p < 0.01. **(E)** Active αM (mAb24 high-affinity conformation) expression was measured by flow cytometry and analyzed as in (D). 0 vs. 1 nM, ns; 0 vs. 10 nM, *p < 0.05; 0 vs. 100 nM, **p < 0.01.

### α7nAChR activation reduces neutrophil recruitment during endotoxemia, which has a protective effect on mouse survival

To determine whether the inhibition of neutrophil migration observed ex vivo is recapitulated in vivo, we performed adoptive transfer experiments in a mouse model of LPS-induced endotoxemia. Neutrophils were isolated from mouse bone marrow by negative selection using magnetic beads. The purity of the isolated neutrophil population was assessed by flow cytometry using anti-Ly6G and anti-CD11b antibodies, confirming a highly enriched neutrophil preparation (Fig. 7 B). Neutrophils were divided into two groups: one group was labeled with PKH67 green fluorescent dye and pretreated with PNU-282987 for 30 min, whereas the second group was labeled with PKH26 red fluorescent dye and treated with vehicle. Equal numbers of green- and red-labeled neutrophils (2 × 10⁶ total cells) were mixed and injected intravenously into recipient mice immediately before intraperitoneal administration of LPS. Therefore, equal numbers of both populations were co-transferred into the same recipient mouse to directly compare the migratory capacity of PNU-treated and control neutrophils under identical inflammatory conditions. (Fig. 7A).

**Figure 7.**
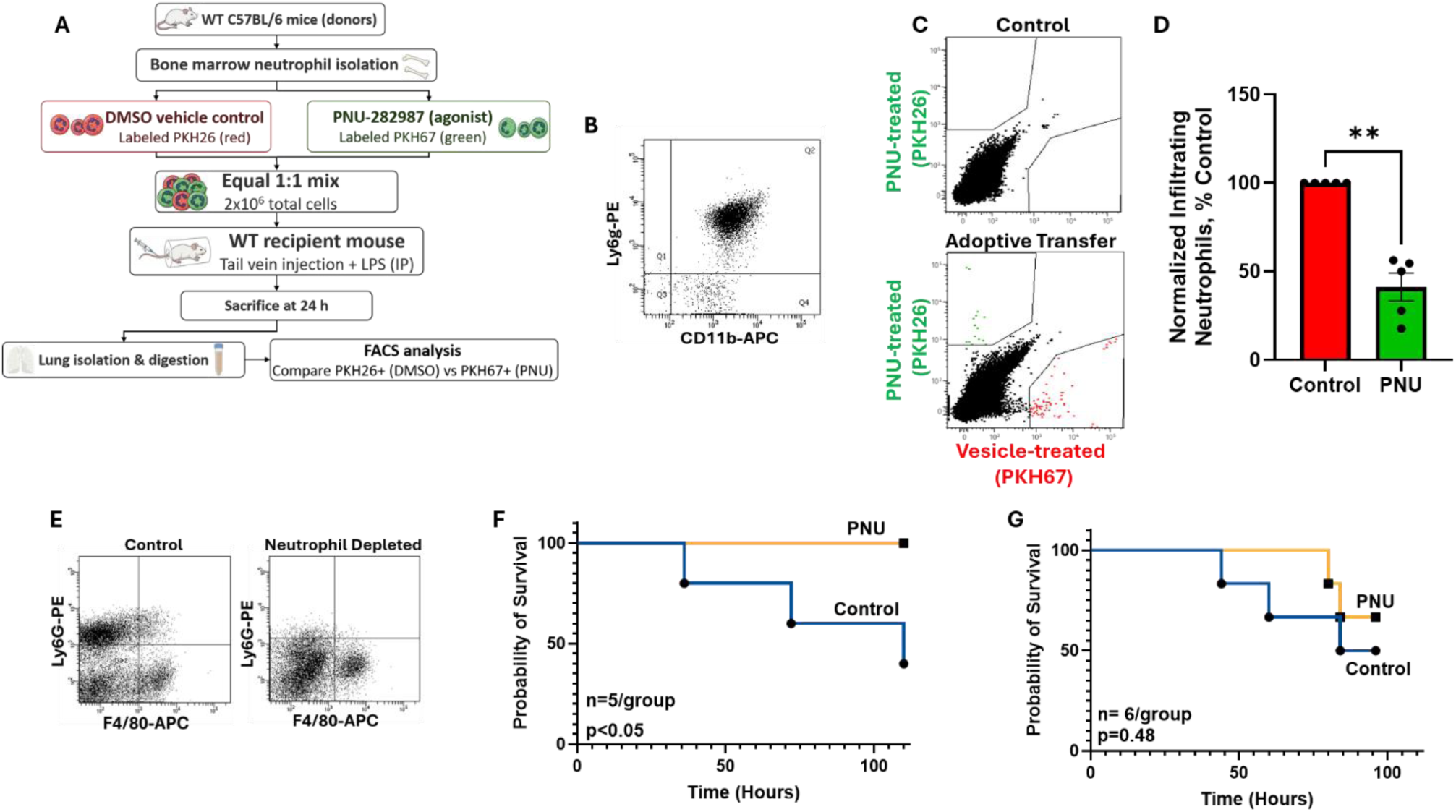
α7nAChR activation attenuates neutrophil lung recruitment and improves survival during LPS-induced endotoxemia in a neutrophil dependent manner. **(A)** Schematic representation of the adoptive transfer experimental workflow: recipient mice received co-transferred fluorescently labeled (PKH67 or PKH26) neutrophils pretreated with vehicle or PNU-282987 (30 µM) prior to LPS challenge. **(B)** Representative flow cytometry dot plot confirming purity of magnetic-bead-isolated bone marrow neutrophils (Ly6G^+^/CD11b^+^) (>90%). **(C)** Flow cytometry analysis of digested lung tissue following transcardial perfusion, gating on transferred PKH-labeled neutrophil populations in the lungs. **(D)** Quantification of relative transferred neutrophil migration into lung tissue. Data from individual experiments were normalized to vehicle control for each experimental day. Data are presented as individual data points with bars representing mean ± SEM (n=5). Statistical significance was determined using a Welch’s t-test, p<0.01. **(E)** Representative flow cytometry validation of anti-Ly6G antibody-mediated neutrophil depletion (Ly6G+ vs. F4/80+) compared to untreated controls. Mice were depleted of neutrophils via IP administration of anti-Ly6G antibody (100 µg). (F) WT C57BL/6J mice (n = 5/group) received PNU-282987 (3 mg/kg, IP) or DMSO vehicle 30 min prior to LPS challenge (8 mg/kg, IP), and survival was monitored for up to 120 h. Survival curves were compared by log-rank (Mantel-Cox) test. **(G)** Mice (n = 6/group) were depleted of neutrophils via IP administration of anti-Ly6G antibody (100 µg) 24 h before and 24 h after LPS challenge, then treated with PNU-282987 or vehicle and challenged with LPS (10 mg/kg, IP). Survival was monitored for up to 96 h and curves were compared by log-rank (Mantel-Cox) test.

Twenty-four hours later, mice were euthanized, perfused, and lungs were harvested for flow cytometric analysis. The analysis demonstrated that PNU-282987-treated neutrophils accumulated in significantly lower numbers in the lungs than vehicle-treated neutrophils (Fig. 7 C, D), indicating that activation of α7nAChR directly suppresses neutrophil recruitment during endotoxemia. Notably, our previous studies demonstrated that exchanging PKH67 and PKH26 labeling between experimental groups produced equivalent migration results, indicating that the observed cell migration was not influenced by the choice of fluorescent dye.^29^ These findings are consistent with our ex vivo swarming assay and with our previous observation that systemic administration of PNU-282987 reduces pulmonary neutrophil infiltration during endotoxemia.^29^

We next investigated whether neutrophils contribute to the protective effects of α7nAChR activation. Neutrophils were depleted using the widely established clone 1A8 antibody, which was administered 24 h before and 24 h after LPS challenge. The efficiency of neutrophil depletion was verified by flow cytometric analysis of enzymatically digested lung tissue collected 24, 48, and 72 h after LPS challenge (Fig 7E). Mice received PNU-282987 (3 mg/kg) 30 min before LPS administration (10 mg/kg), and survival was monitored for 96 h. As expected, PNU-282987 significantly improved survival in the control experiment with wild-type mice (Fig. 7F), consistent with our previous findings^29^. In contrast, neutrophil depletion markedly diminished the protective effect of PNU-282987, resulting in mortality not significantly different from that of vehicle-treated control mice (Fig. 7G). These results demonstrate that neutrophil depletion reduced the defensive effect of PNU-282987 and indicate that, in addition to well-established macrophages, neutrophils may contribute to α7nAChR-mediated protection during endotoxemia.

## DISCUSSION

Here, we identify neutrophils as a direct cellular target of α7nAChR-mediated cholinergic signaling. We demonstrate that α7nAChR activation suppresses neutrophil migration by limiting α_M_β_2_ activation, actin remodeling, and recruitment during endotoxemia. These findings provide novel insights into the cholinergic anti-inflammatory pathway, previously associated with macrophage targeting, and establish neutrophils as active contributors to its protective effects.

The role of macrophages in mediating the cholinergic anti-inflammatory pathway is well documented, including the inhibition of NF-κB signaling^13, 43, 44^ and the regulation of macrophage migration^29^. In contrast, the role of neutrophils remained poorly understood. While cholinergic stimulation reportedly reduces neutrophil accumulation at sites of inflammation^9^, the underlying mechanisms remained largely unknown ^9, 14, 15^. The most detailed study suggested that α7nAChR expressed on endothelial cells transmits signals that suppress neutrophil migration^45^. Here, we demonstrate that cholinergic stimulation acts directly on neutrophils through α7nAChR. The receptor specificity of this response is supported by the use of two structurally distinct α7nAChR agonists and by the loss of the inhibitory effect in α7nAChR-deficient neutrophils.

We demonstrate an important role for integrin α_M_β_2_ in the cholinergic regulation of neutrophil migration. Our data show that α7nAChR-mediated signaling reduces α_M_β_2_-mediated adhesion, thereby altering neutrophil migratory behavior. Integrin α_M_β_2_ is expressed on resting neutrophils in a bent, low-affinity conformation^46^. Inflammatory stimulation induces conformational changes that shift the integrin into an intermediate activation state, allowing ligand binding with intermediate affinity^47^. Subsequent inside-out signaling mediated by G-protein-coupled receptor agonists such as fMLP, LTB4, or TNFα converts α_M_β_2_ into a fully active, high-affinity conformation that promotes robust cell adhesion and migration^26^. However, inflammatory stimulation also induces the mobilization of intracellular α_M_β_2_-containing granules, increasing surface α_M_β_2_ expression by approximately 3- to 5-fold within 15 min^23, 24^. Elevated integrin density together with high integrin affinity may generate excessive adhesive forces that impair neutrophil migration. Therefore, depending on the degree of activation, inflammatory stimulation can either promote or inhibit neutrophil migration.

This concept is consistent with the well-established biphasic relationship between integrin-mediated adhesion and cell migration. Cell motility follows a bell-shaped curve in response to substrate adhesiveness, with maximal migration occurring at intermediate adhesive strength. Insufficient adhesion prevents efficient traction, whereas excessive adhesion causes cell arrest. Ligand density, integrin affinity, and integrin surface expression collectively determine this balance ^48–50^.

Results from the swarming experiments using the β_2_-integrin activating antibody MEM48 or fMLP further support the concentration-dependent role of α_M_β_2_ activation in neutrophil migration and explain the differential effects of PNU-282987. PNU-282987 reduced swarming in both untreated and MEM48-treated neutrophils. However, although PNU decreased α_M_β_2_ activation under both conditions, the biological consequences varied because the initial levels of integrin activation were markedly different. Untreated neutrophils are physiologically activated by gradients of fMLP and LTB4, which generate an optimal level of adhesion for migration. By reducing α_M_β_2_ activation below this optimum, PNU-282987 decreases adhesion and consequently inhibits migration. In contrast, MEM48 induces pseudo-physiological α_M_β_2_ activation and excessive adhesion. Under these conditions, PNU-282987 shifts α_M_β_2_ toward an intermediate activation state, reducing excessive adhesion and partially restoring neutrophil migration.

Although MEM48 activates all β_2_ integrins, including α_M_β_2_, α_L_β_2_, α_X_β_2_, and α_D_β_2_, our data indicate that α_M_β_2_ is the principal receptor responsible for the observed effects. First, α_L_β_2_ does not bind fibrinogen and therefore does not contribute to fibrinogen-dependent migration^51^. Second, α_X_β_2_ and α_D_β_2_ are expressed at substantially lower levels than α_M_β_2_ on neutrophils, making their contribution to fibrinogen-dependent migration likely minimal^37, 52^.

Our data show the marked difference in F-actin polymerization during swarming of control and PNU-treated neutrophils. The characteristic actin cup on the leading edge and uropod tail was lost in α7nAChR-activated neutrophils, providing a structural explanation for their impaired migration. Notably, it has been suggested previously that actin polymerization is affected in neutrophils of nicotine-treated mice^45^. This finding was attributed to a direct effect of α7nAChR signaling on the actin cytoskeleton, resulting in reduced integrin expression at the cell surface. In contrast, our data indicate that integrin α_M_β_2_ functions upstream of actin polymerization in the α7nAChR-mediated signaling pathway. Importantly, the effect of cholinergic stimulation on actin remodeling was abolished in neutrophils isolated from α_M_-deficient mice. In our model, PNU treatment reduced the activation of α_M_β_2_ that prevents clustering, integrin-mediated signaling and formation of actin cytoskeleton, which is required for proper cell migration. Together, these findings identify α_M_β_2_ as the key upstream regulator between α7nAChR signaling and cytoskeletal remodeling.

Interestingly, α7nAChR activation exerts opposite effects on macrophage and neutrophil migration. Whereas our previous work demonstrated that α7 signaling promotes macrophage recruitment during endotoxemia^29^, the present study shows that the same pathway suppresses neutrophil migration. These divergent responses likely reflect fundamental differences in integrin utilization and adhesion requirements between the two leukocyte populations^40, 41^.

Although our data identify α_M_β_2_ as a major downstream effector of α7nAChR signaling, the intracellular signaling events linking receptor activation to integrin deactivation remain to be defined. Likewise, additional studies will be required to determine whether similar mechanisms regulate neutrophil trafficking in infectious sepsis and other inflammatory diseases.

## CONCLUSIONS

Our findings identify neutrophils as previously unrecognized effectors of the cholinergic anti-inflammatory pathway and establish α_M_β_2_ as a key molecular effector through which α7nAChR regulates neutrophil migration. These observations expand current understanding of neuroimmune regulation and suggest that selective modulation of α7nAChR–α_M_β_2_ signaling may be a promising therapeutic strategy for inflammatory disorders.

## Abbreviations

α7nAChR: alpha 7 nicotinic acetylcholine receptor
WT: wild-type
LPS: lipopolysaccharide
BSA: bovine serum albumin
PBS: Phosphate-buffered saline
DMSO: Dimethyl Sulfoxide
EDTA: Ethylenediaminetetraacetic acid
HBSS: Hank’s Balanced Salt Solution
MACS: Magnetic-activated Cell Sorting
PFA: Paraformaldehyde
IP: Intraperitoneal
LTB4: Leukotriene B4
PMA: Phorbol 12-myristate 13-acetate
TNFα: Tumor Necrosis Factor alpha
IL-1β: Interleukin-1β
IL-6: Interleukin-6

## Declarations

### Ethics approval and consent to participate

The collection and isolation of human neutrophils were conducted in strict accordance with federal regulations and institutional guidelines, as previously described. All experimental protocols were reviewed and approved by the Institutional Biosafety and Chemical Safety Committee at East Tennessee State University. Fresh peripheral blood samples from healthy donors were purchased from a local BioIVT collection center. Blood was collected into EDTA tubes under protocols approved by the BioIVT Institutional Review Board, and all samples were de-identified before being provided for research use.

All animal procedures were performed according to animal protocols approved by East Tennessee State University IACUC.

### Consent for Publication

Not applicable.

### Availability of Data and Materials

The datasets used and/or analyzed during the current study are available from the corresponding author on reasonable request.

### Authors’ Contributions

JLC designed and performed the experiments, analyzed the data, and wrote the manuscript. TP, SS, TT, and AT performed experiments and collected resulting data. AH and VAP designed the research, analyzed the data and edited the manuscript. VPY conceptualization, designed the research, analyzed the data, and wrote the manuscript. All authors read and approved the final manuscript.

## Acknowledgements

Not applicable.

